# Population structure and antimicrobial resistance of *Corynebacterium diphtheriae* in Victoria, Australia

**DOI:** 10.1101/2025.07.23.666245

**Authors:** Lamali Sadeesh Kumar, Kylie Hui, Janet Strachan, Norelle L. Sherry, Benjamin P. Howden, Sarah L. Baines

## Abstract

*Corynebacterium diphtheriae*, the main aetiological agent of diphtheria, is a re-emerging bacterial pathogen of public health concern, yet remains largely understudied globally. In this study, we analysed the population structure and antimicrobial resistance (AMR) of 210 *C. diphtheriae* isolates from Victoria, Australia including 103 historical (1950–1970) and 107 contemporary clinical isolates (2004–2023), using whole-genome sequencing and phenotypic susceptibility testing. The diphtheria toxin gene (*tox*) was detected in 89 isolates, the majority of which (n=83; 93.3%) were historical. Population structure comprised two primary phylogenetic lineages, Mitis and Gravis, each containing multiple sublineages. Multi-locus sequence type (MLST) analysis revealed a highly diverse population structure with multiple novel MLST profiles and alleles. When placed within a global phylogenetic framework, Australian isolates were broadly distributed, reflecting substantial genetic diversity. Phenotypic susceptibility testing against eleven antimicrobials revealed that several contemporary isolates were resistant to multiple agents, including penicillin and erythromycin, first-line treatments of *Corynebacterium* infections. Eight contemporary isolates were multidrug-resistant (resistant to ≥3 antimicrobial classes), including five with resistance to both penicillin and erythromycin. Genomic analysis identified multiple genes and mutations conferring resistance among contemporary isolates. In contrast, no antimicrobial resistance phenotypes or genotypes were observed in historical isolates. Analysis of historical genomes provides valuable insights into a period of heightened diphtheria activity in Victoria prior to widespread immunisation. Overall, these findings establish a baseline for ongoing genomic surveillance in the face of increasing global outbreaks, support informed empiric treatment strategies, and contribute to the knowledge of global population structure of *C. diphtheriae*.

**Impact statement:** *Corynebacterium diphtheriae* is a re-emerging pathogen of public health concern with limited genomic data available to support surveillance and public health interventions. This study provides a contemporary understanding of *C. diphtheriae* from Australia in the era of genomic surveillance and helps us better understand baseline genomic diversity and antimicrobial resistance. Additionally, these findings enhance preparedness for potential future incursions or outbreaks, including those involving drug-resistant strains, as recently observed in parts of Africa and Europe.

**Data summary:** Genome sequences are deposited in GenBank under BioProject PRJNA870170. Sample data and accession numbers are included in the Supplementary Table S1. The authors confirm all supporting data, code and protocols have been provided within the article or through supplementary data files.

## Introduction

*Corynebacterium diphtheriae* is a Gram-positive rod and the main aetiological agent of diphtheria, when toxigenic. Toxigenicity occurs when the *C. diphtheriae* (or other related *Corynebacterium* species) is lysogenised by corynbacteriophages that carry the diphtheria toxin gene (*tox*) [1].

Diphtheria is a vaccine-preventable disease that can cause respiratory or cutaneous illness, with transmission occurring via respiratory droplets or through contact with infected ulcers. It was a major public concern in Australia with high morbidity and mortality rates in early 1900s prior to the introduction of diphtheria toxoid vaccine. However, nationwide immunisation programs have resulted in a dramatic decline of diphtheria cases in Australia and many other countries. [2, 3]. Nowadays, diphtheria is rarely reported in Australia, with a small number of sporadic cases, mostly acquired from outside of Australia [4].

Immunisation against diphtheria provides protection only against toxigenic *C. diphtheriae* strains. However, multiple illnesses including invasive diseases such as bacteraemia, endocarditis, and septic arthritis caused by non-toxigenic strains have been reported worldwide [5–8], with increasing incidence of disease caused by non-toxigenic strains reported from multiple world regions [9–11].

Widespread use of improved bacterial identification methods such as matrix assisted laser desorption-ionization time-of-flight mass spectrometry (MALDI-TOF MS) systems have enhanced the detection of *tox* negative strains in the last decade [12, 13]. In addition to toxigenic and non-toxigenic strains, non-toxigenic *tox* gene-bearing (NTTB) *C. diphtheriae* strains have also been reported from multiple countries including Australia [14]. By definition, NTTB strains are genetically positive for the *tox* gene, yet phenotypically negative for diphtheria toxin production, and considered as possible reservoirs for the *tox* gene in populations. While inactivating mutations in the *tox* gene may explain the lack of toxin production in some NTTB strains, no definitive genetic explanation has been determined in others. [15, 16].

The primary treatment for diphtheria (other than supportive measures) is administration of diphtheria antitoxin (DAT) to neutralise unbound toxin and reduce adverse systemic effects of diphtheria toxin. Antimicrobials are prescribed for all *C. diphtheriae* infections regardless of toxigenicity and recommended first-line antimicrobials include penicillin and erythromycin. Additionally, close contacts of a patient with confirmed diphtheria are prescribed antimicrobial therapy as chemoprophylaxis, along with primary or booster vaccination against diphtheria if necessary [1, 16, 17]. However, reduced susceptibility or resistance to first-line antimicrobials and other antimicrobial agents, including multidrug resistance, have been reported globally [16, 18–22]. Multiple genes and mutations have been described in *C. diphtheriae* conferring antimicrobial resistance (AMR), including the *pbp2m* gene [16] and *erm(X)* gene [23] conferring resistance to penicillin and erythromycin accordingly.

Recently, *C. diphtheriae* has gained attention as a re-emerging pathogen causing multiple diphtheria outbreaks globally [22, 24], including strains noted to exhibit significant AMR. Although toxigenic *C. diphtheriae* strains remain rare in Australia, non-toxigenic strains are regularly identified [4, 8, 13]. Thus, it is important to establish and maintain surveillance of *C. diphtheriae* strains in Australia for outbreak detection and response, and to monitor AMR.

Here we present a genomic portrait of *C. diphtheriae* recovered from the state of Victoria, Australia, over contemporary (2004-2023) and historical (1950-1970) years. This study aims to provide critical insights into the prevalence of the diphtheria toxin gene, phenotypic antimicrobial susceptibility profiles, and associated genetic determinants of AMR, as well as to investigate temporal trends in these clinically relevant features. Additionally, we aim to establish a phylogenetic framework for understanding the local population structure of Victorian *C. diphtheriae* within a global context, thereby enhancing our understanding of the pathogen’s epidemiology in preparation for future incursions and outbreaks.

## Methods

### Study design and dataset

The study was performed at the Microbiological Diagnostic Unit Public Health Laboratory (MDU PHL) in the state of Victoria, Australia (population 7.0 million at 30 September 2024) [25], the laboratory responsible for statewide surveillance of bacterial public health pathogens for over 125 years, with an extensive collection of historical isolates. A total of 210 *Corynebacterium diphtheriae* isolates was included in this study (Supplementary Table S1). The dataset was divided into two subsets; contemporary clinical isolates (n=107) submitted by diagnostic laboratories in Victoria between 1st January 2004 and 31st December 2023, and a selection of historical isolates (n=103) submitted between 1950 and 1970, revived from vials of freeze-dried material. Historical isolates were chosen to represent a range of years from 1950-1970. Duplicate isolates were identified and excluded, with the earliest isolate for each patient being included in the study. All isolates were identified as *C. diphtheriae* by VITEK® MS MALDI-TOF technology.

### DNA extraction, whole genome sequencing and genome analysis

*C. diphtheriae* cultures were grown aerobically at 37°C for 24h on horse blood agar. Genomic DNA extraction was performed using the Virus/Pathogen DSP mini kit on the QIAsymphony (QIAGEN). DNA libraries were prepared using the Nextera XT kit following manufacturer’s instructions (Illumina). Whole genome sequencing was performed on the Illumina NextSeq 500/550 systems using 2 x 150 bp paired-end chemistry following manufacturer’s instructions.

The Bohra bioinformatics pipeline (v2.3.7) (https://github.com/MDU-PHL/bohra) was used for quality control of sequence reads and genome assembly. SKESA (v2.5.1) (https://github.com/ncbi/SKESA) [26] was used as the *de novo* sequence assembler, run with default parameters. *In silico* multi-locus sequence typing (MLST) was performed with mlst (v2.23.0) (https://github.com/tseemann/mlst) following the MLST scheme published by the Institut Pasteur (https://bigsdb.pasteur.fr/diphtheria/). *C. belfantii* and *C. rouxii* strains, formerly classified as *C. diphtheriae*, were identified by genome-wide average nucleotide identity (ANI) on genomic sequence data as previously described [27], and were subsequently excluded from the study.

### Detection of diphtheria toxin gene (*tox*)

All assembled sequences were screened for diphtheria toxin (*tox*) gene with AMRFinderPlus (v3.12.8). Additionally, *tox* gene detection for the subset of contemporary isolates was performed routinely at MDU PHL via a conventional PCR assay at the time of sample receipt [28]. Aligned *tox* gene sequences from assemblies against the *tox* gene sequence from the reference strain NCTC 13129 (NC_002935.2) were obtained using gene-puller (identity >95% and coverage >90%) and visualised with Geneious Prime (Version 2024.0.7) (http://www.geneious.com) to investigate any genotypes related to NTTB strains as described previously [15, 16].

### Victorian phylogenetic structure of *C. diphtheriae*

A whole genome alignment of Victorian *C. diphtheriae* isolates was produced with snippy (v4.6.0) (https://github.com/tseemann/snippy) using *C. diphtheriae* NCTC 13129 (NC_002935.2) as the reference genome. A maximum likelihood (ML) tree was inferred with IQ-TREE 2 [29] using the best-fit model TVM+F+R10. Effects of homologous recombination events were quantified and masked with ClonalFrameML (v1.13) [30]. The ML phylogenetic tree was visualized and annotated with Interactive Tree of Life (iTOL) (v6) [31].

Previous studies have defined two main phylogenetic lineages, Mitis and Gravis, after observing a strong correlation with presence (Gravis) or absence (Mitis) of the *spuA* gene [16]. Hence, the presence of *spuA* was investigated in genome assemblies using fastablasta (identity >90% and coverage >95%) (https://github.com/kwongj/fastablasta).

### Global phylogenetic structure of *C. diphtheriae*

The global population structure of *C. diphtheriae* was analysed using all 210 strains from this study alongside 885 publicly available strains, representing six continents. (Supplementary Table S2). Only isolates derived from human clinical specimens were selected for analysis. Of the 1,095 isolates, those collected between 2004 and 2023 were classified as contemporary (n= 823), while those collected prior to 2004 were classified as historical (n= 62). A phylogenetic tree for global population structure was created using mashtree (https://github.com/lskatz/mashtree) (--sketch-size 50000 --seed 100 --min-depth 0). The mashtree was visualized and annotated with Interactive Tree of Life (iTOL) (v6) [31].

### Antimicrobial Susceptibility Testing

Phenotypic antimicrobial susceptibility testing (AST) was performed by broth microdilution (BMD) for the following 11 antimicrobials: ciprofloxacin, clindamycin, erythromycin, gentamicin, linezolid, penicillin, quinupristin/dalfopristin, rifampicin, tetracycline, trimethoprim/sulfamethoxazole, and vancomycin. BMD was performed with direct colony suspensions equivalent to a 0.5 McFarland standard incubated in cation-adjusted Mueller-Hinton broth with lysed horse blood (2.5% to 5% v/v) at 35°C in ambient air for 24hrs. Sensititre AIM™ Automated Inoculation Delivery System was used for inoculating the broth to GPN3F Sensititre standard plates (Thermo Fisher Scientific, Waltham, MA, USA). Following incubation, minimum inhibitory concentration (MIC) values were read using the Sensititre ARIS HiQ System or Sensititre Vizion Digital MIC Viewing System (Thermo Fisher Scientific, Waltham, MA, USA). Quality control was performed with the *Streptococcus pneumoniae* ATCC® 49619 reference strain. Minimum inhibitory concentration (MIC) values for the above antimicrobials were interpreted following the breakpoints for *Corynebacterium* spp. from Clinical and Laboratory Standards Institute (CLSI) M45 guidelines, 3^rd^ ed. [32]. *C. diphtheriae* strains that exhibited resistance to three or more antimicrobial classes were categorised as multidrug-resistant (MDR) *C. diphtheriae*.

### Detection of antimicrobial resistance determinants

Known AMR mechanisms including acquired AMR genes and *C. diphtheriae* specific mutational resistance were identified by analysing the genomic assemblies using AMRFinderPlus (v3.12.8) with *Corynebacterium diphtheriae* as the species option. AMR genes with >90% coverage and >95% identity compared to the AMRFinderPlus database were selected for analysis.

## Results

### Characteristics of *C. diphtheriae* isolates in Victoria

All 210 isolates of this study were confirmed as *Corynebacterium diphtheriae* with ANI values above 97% when compared with *C. diphtheriae* type strain NCTC11397^T^, thereby excluding *C. belfantii* and *C. rouxii.* Contemporary *C. diphtheriae* isolates (n=107) were collected from a range of clinical sample types, including cutaneous (n=77), respiratory (n=14), blood (n=3), other sources such as tissue, pus, and ear swabs (n=8), and five isolates of unknown source (Supplementary Table S1). No metadata were available for historical isolates (n=103).

A total of 89 (42.4%) *C. diphtheriae* genomes were found to carry the *tox* gene, while the remaining 121 (57.6%) isolates were *tox* negative. The *tox* gene was detected in 80.6% (83/103) of historical isolates and only 5.6% (6/107) of contemporary isolates. PCR and *in silico* genomic detection of *tox* gene were 100% concordant for *tox* positive contemporary isolates. Among all *tox* positive isolates, the *tox* gene remained intact, with no internal stop codons or other disruptive genetic events observed, such as those reported in NTTB strains.

For *tox* positive historical isolates (n=83), available metadata were insufficient to assess potential epidemiological links between cases. The *tox* positive contemporary isolates were reported in 2013 (n=2), 2017 (n=2), 2019 (n=1), and 2022 (n=1), with no identified epidemiological links between the cases. However, all six patients had a recent history of overseas travel, suggesting that these strains were likely imported (Supplementary Table S1).

### Victorian *C. diphtheriae* isolates were highly diverse

MLST analysis revealed significant diversity, with 50 different known sequence types (STs) identified across historical and contemporary isolates. Novel MLST profiles and/or allelic sequences were observed in a total of 79 isolates, from both contemporary (n=22) and historical (n=57) subsets. ST32 was the most frequently reported ST among contemporary isolates (n=17), and it was exclusive to the contemporary subset. All isolates of ST32 were *tox* negative and detected across multiple years between 2008 and 2023. In comparison, historical isolates showed a higher prevalence of ST25 (n=13), and ST67 (n=14), all of which were *tox* positive (Supplementary Table S1).

Only three STs, ST67, ST124, and ST20, were observed in both historical and contemporary subsets. Isolates of ST67 were *tox* positive in both historical and contemporary subsets. In contrast, historical isolates of ST124 and ST20 were all *tox* positive while contemporary isolates of the same STs were *tox* negative (Supplementary Table S1).

Phylogeny of *C. diphtheriae* strains from Victoria rooted with *C. belfantii* exhibited a star-like phylogeny with multiple deep branching clades. As per ClonalFrameML (v1.13) results, the relative rate of recombination to mutation (R/theta) was 0.64. Average length of recombination segments (delta) was 599 bp and the mean genetic distance between donor and recipient of recombination (nu) was 0.02. This resulted in a relative impact of recombination to mutation (r/m = R/theta × delta × nu) of 7.0, which means that the homologous recombination impacts the *C. diphtheriae* diversification seven times more than mutations.

The distribution of the *spuA* gene largely distinguished the two previously described phylogenetic lineages, Mitis and Gravis. Both lineages were equally represented among Victorian *C. diphtheriae* isolates. All but 10 Gravis lineage isolates were *spuA*-positive, and all but two Mitis lineage isolates were *spuA*-negative. Both Mitis and Gravis lineages were represented in both historical and contemporary isolates. *tox* positive and negative isolates were found in both the Mitis and Gravis lineages; however, a higher proportion of *tox* positive isolates was observed in the Mitis lineage (57/115; 50%) compared to the Gravis lineage (32/95; 33.7%) (Supplementary Table S1, Figure 1).

**Figure 1:**
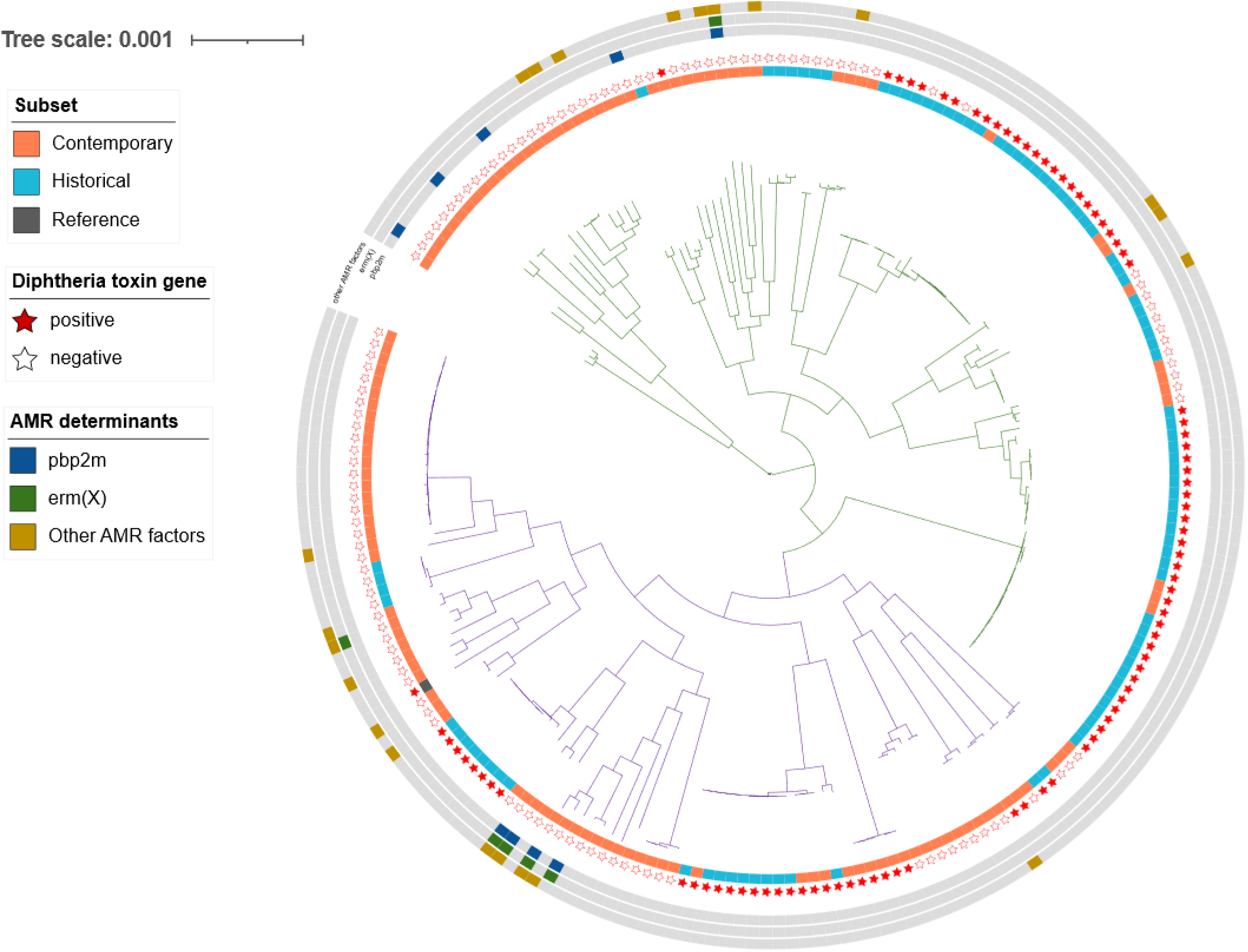
Phylogenetic tree (maximum likelihood-recombination adjusted) of 210 *Corynebacterium diphtheriae* isolates from Victoria, Australia. The tree is constructed from core genome SNPs using *C. diphtheriae* NCTC13129 as the reference. Tree is rooted at *C. belfantii* (FRC0043a) and the branches of the main phylogenetic lineages, Mitis and Gravis are coloured in green and purple respectively. The inner ring corresponds to contemporary (*n*=107, orange) and historical (*n*=103, blue) isolates, and reference strain, *C. diphtheriae* NCTC 13129 (grey). The outer ring interprets the presence (filled star) or absence (unfilled star) of diphtheria toxin gene (*tox*) status. The three outer circles represent the presence of genetic determinants of antimicrobial resistance (AMR): *pbp2m* (dark blue), *erm(X)* (dark green) and other AMR factors (dark yellow).

### Australian isolates were widely dispersed throughout the global phylogeny

Analysis of the global phylogenetic structure of *C. diphtheriae* revealed that isolates from Victoria, Australia (this study) were highly diverse and dispersed across the global phylogeny (Figure 2). *C. diphtheriae* isolates from other Australian studies were also distributed across the global phylogeny but appeared to be clustered closely to the Victorian isolates, indicating that Australian *C. diphtheriae* isolates of the same clade appear to have national distribution and were not unique to smaller geographic regions (states/territories) (Figure 2).

**Figure 2:**
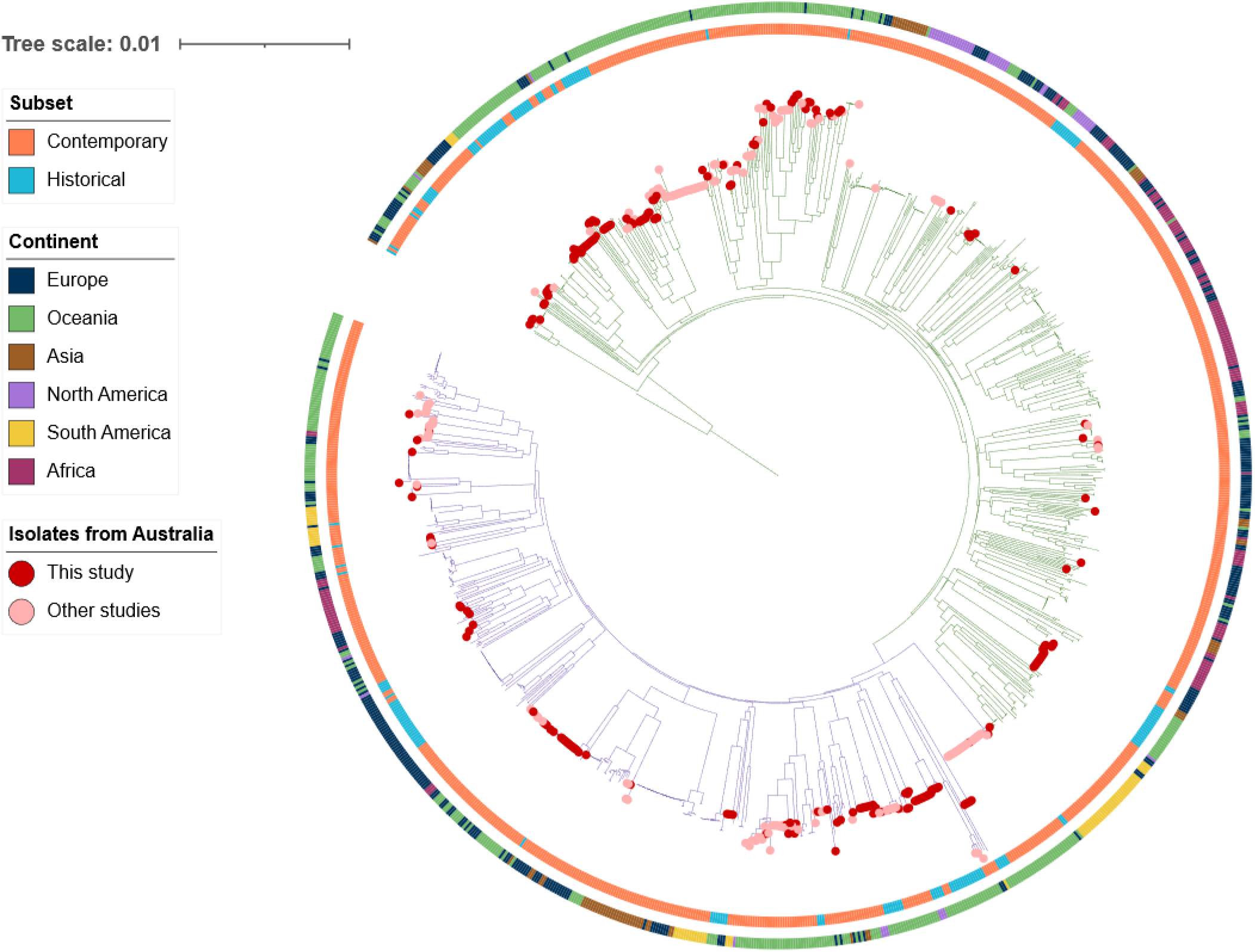
Phylogenetic tree (mashtree) of 1,095 *Corynebacterium diphtheriae* isolates from six continents. The tree is rooted at *C. diphtheriae* SAMN13343853, and the branches of the main phylogenetic lineages, Mitis and Gravis are coloured in green and purple respectively. The inner ring around the tree corresponds to contemporary isolates (2004-2023, orange) and historical isolates (prior to 2004, blue). The outer ring is coloured by continent where the isolate was reported. Symbols at the branch tips represent Australian *C. diphtheriae* isolates (n=384); Victorian isolates from this study (red) (n=210) and Australian isolates from other studies (pink) (n=174).

### Phenotypic antimicrobial resistance was only present in contemporary strains

Antimicrobial susceptibility testing of all 210 *C. diphtheriae* isolates revealed susceptibility profiles for 11 antimicrobials, including the first-line antimicrobials: penicillin and erythromycin. All historical isolates (n=103) were susceptible to 10/11 antimicrobial agents: ciprofloxacin, clindamycin, erythromycin, gentamicin, linezolid, quinupristin/dalfopristin, rifampicin, tetracycline, trimethoprim/sulfamethoxazole, and vancomycin. The majority of historical isolates (n=76, 73.8%) were susceptible to penicillin, while a quarter of the subset (n=27, 26.2%) tested as intermediate to penicillin, as per CLSI breakpoints [32] for *Corynebacterium* spp. No penicillin resistant phenotypes were observed in historical isolates (Figure 3, Supplementary Table S3).

**Figure 3:**
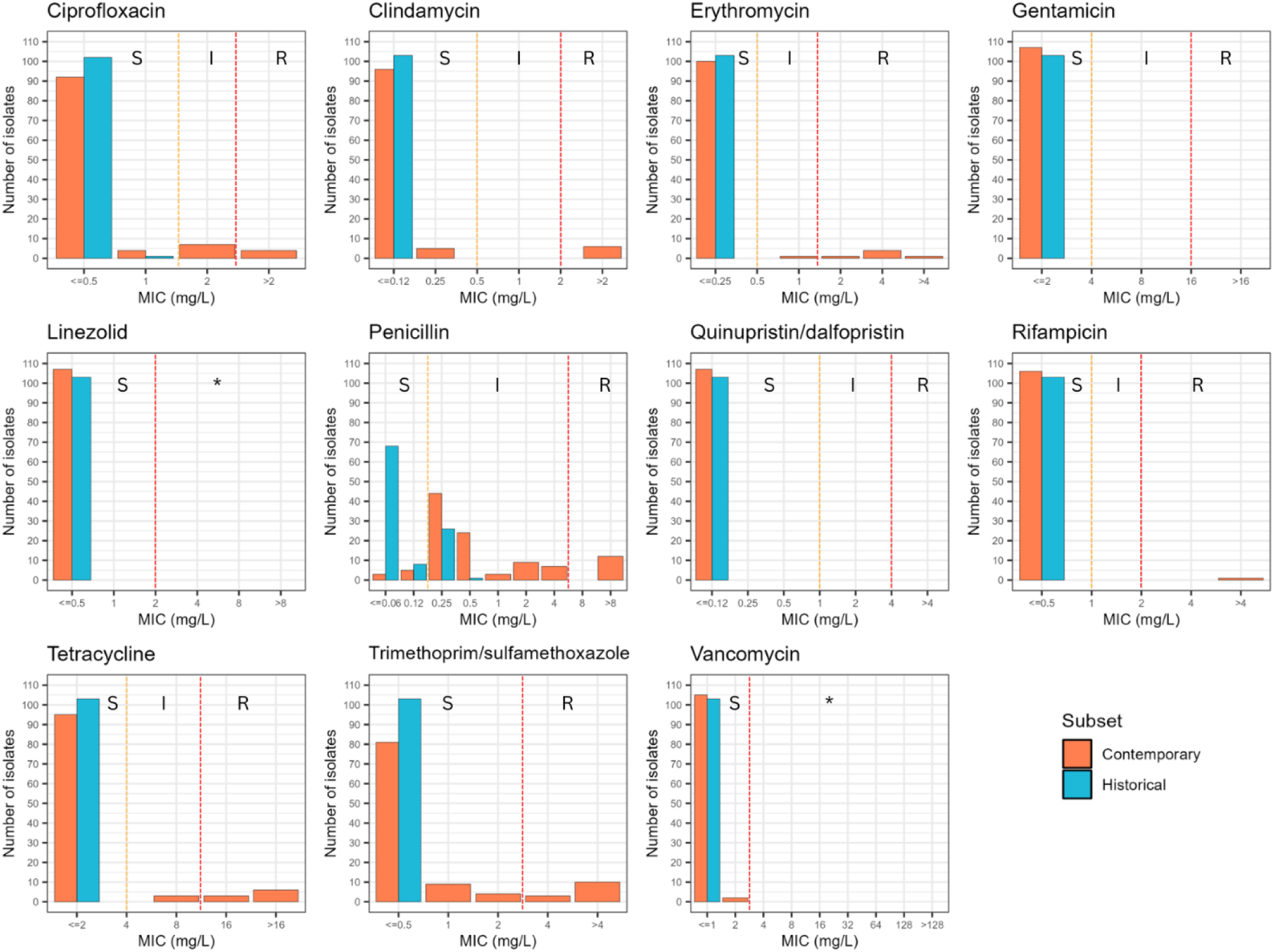
Antimicrobial susceptibility of *C. diphtheriae* isolates. Distribution of minimum inhibitory concentration (MIC) values of 11 antimicrobials obtained from broth microdilution for contemporary (n=107) and historical (n=103) isolates Colour of the bars represents contemporary (orange) and historical (blue) isolates of *C. diphtheriae*. MIC values were interpreted as susceptible (S), intermediate (I) or resistant (R) following the Clinical and Laboratory Standards Institute (CLSI) M45 (3^rd^ ed.) guidelines. Asterisk (*) denotes ‘Non-susceptible’ category as per CLSI M45 guidelines (2015). Absence or rare occurrence of resistant strains is reported for this organism/antimicrobial agent combination.

All contemporary isolates (n=107) were susceptible to 4/11 antimicrobial agents: gentamicin, linezolid, quinupristin/dalfopristin, and vancomycin. A small number of contemporary isolates (7.5%, n=8) were fully susceptible to all 11 antimicrobial agents. Resistant phenotypes were observed for 7/11 antimicrobial agents: ciprofloxacin, clindamycin, erythromycin, penicillin, rifampicin, trimethoprim/sulfamethoxazole and tetracycline (Figure 3, Supplementary Table S3). Phenotypic resistance to at least one antimicrobial agent was observed in 30.8% (n=33) of contemporary isolates. For penicillin, only 7.5% (n=8) of contemporary isolates tested as susceptible while 74.8% (n=80) and 17.7% (n=19) tested as intermediate and resistant respectively. For erythromycin, 93.4% (n=100) contemporary isolates tested as susceptible, while 0.9% (n=1) and 5.6% (n=6) tested as intermediate and resistant respectively (Supplementary Table S3).

Notably, eight contemporary isolates were identified as multidrug-resistant (MDR) *C. diphtheriae*. Five MDR isolates were resistant to both first-line antimicrobials: penicillin and erythromycin while the remaining three isolates were not resistant to either of first-line antimicrobials. All MDR isolates were non-toxigenic isolates from cutaneous samples, isolated between 2016-2023. Multidrug resistance was not observed among *tox* positive isolates. However, two *tox* positive contemporary isolates (of ST243); were resistant to tetracycline, and one was further resistant to trimethoprim/sulfamethoxazole (Supplementary Table S3).

### Prevalence and associations of genetic AMR determinants

A search for AMR genotypes revealed that there were no known AMR genes or mutations present in the historical *C. diphtheriae* isolates. However, 27 contemporary isolates carried at least one previously reported AMR determinant. These included antimicrobial genes conferring resistance to beta-lactams (*pbp2m*), macrolides (*erm(X)*), aminoglycosides (*aph(3’)-Ia*, *aph(6)-Id*, and *aph(3’’)-Ib*), tetracyclines (*tet(W)*, *tet(33)*, and *tet(O)*), phenicols (*cmx*), sulfonamides (*sul1*), trimethoprim (*dfrA15*) as well as *C. diphtheriae-*specific point mutations in *gyrA* and *rpoB* conferring resistance to quinolones (*gyrA*_D93Y, *gyrA*_D93A, *gyrA*_S89F, and *gyrA*_S89V) and rifamycin (*rpoB*_S442F) respectively (Figure 4, Supplementary Table S3). A *bla*_OXA_ gene was observed in a single isolate but had an internal stop codon. Isolates carrying AMR genes or mutations were distributed across the Victorian *C. diphtheriae* phylogeny (Figure 1).

**Figure 4:**
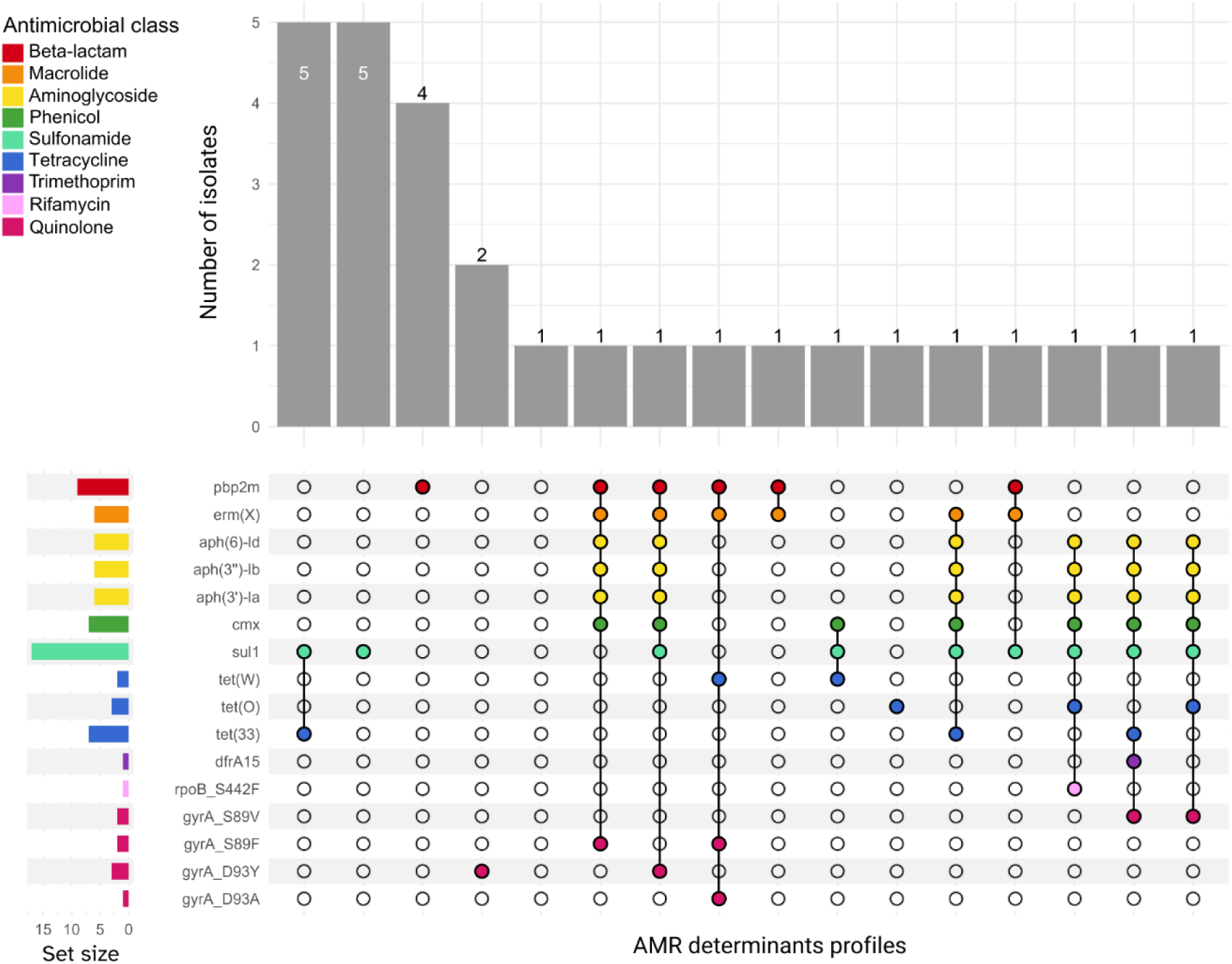
Upset plot of 27 contemporary *Corynebacterium diphtheriae* isolates exhibiting different profiles of genetic antimicrobial resistance (AMR) determinants. Genetic AMR determinants listed in the y-axis were detected by AMRFinderPlus (v3.12.8). The y-axis bars (set-size) demonstrate the total number of isolates with a particular AMR gene or mutation. The strip colours represent the antimicrobial class that the respective gene/ mutation conferring resistance to as listed in the key. X-axis bars represent number of isolates with a particular combination of AMR determinant/s.

A *pbp2m* gene was observed in nine isolates (all tested resistant or intermediate to penicillin) while *erm(X)* was observed in six isolates (all tested as resistant or intermediate to erythromycin). All isolates that tested as resistant or intermediate to tetracycline (n=12) carried one of the three identified tetracycline resistance genes (*tet(W)*, *tet(33)*, or *tet(O)*). All three genes conferring resistance to aminoglycosides (*aph(3’)-Ia*, *aph(6)-Id*, *aph(3’’)-Ib*) were always present together in six contemporary isolates. Phenotypically MDR isolates (n=8) were all *tox* negative and carried a minimum of two and a maximum of eight known AMR determinants. All five MDR isolates with resistance to penicillin and erythromycin carried both *pbp2m* and *erm(X)* genes (Figure 4, Supplementary Table S3).

AMR genes and mutations were rarely observed among *tox* positive isolates, with only two *tox* positive isolates (of ST243) carrying both *sul1* and *tet(O)* genes, consistent with their observed phenotypes (Supplementary Table S3).

## Discussion

Genomic surveillance can provide a comprehensive understanding of a pathogen’s epidemiology and is increasingly being adopted for public health surveillance. In the context of a resurgence of *C. diphtheriae* globally, with the additional threat of AMR complicating treatment and containment [18, 19, 21, 33], we sought to establish the basis for genomic epidemiologic surveillance for public health purposes in our location, where there has been a paucity of genomic data to date. Genomics-enabled public health surveillance of *C. diphtheriae* can support enhanced detection of transmission and monitoring trends for relevant pathogen characteristics, including *tox* carriage and antimicrobial resistance. This study investigated the population structure, *tox* gene carriage and antimicrobial resistance of 210 contemporary and historical isolates of *C. diphtheriae* in Victoria, Australia. We identified a highly diverse population structure of *C. diphtheriae*, similar to what has been described in other settings [16, 24, 33]. Also, we observed that the contribution of homologous recombination was seven times higher to *C. diphtheriae* diversification than mutation, consistent with previous studies [16, 24, 34, 35]. Such extensive recombination enables the rapid acquisition of genetic material, including *tox* gene and AMR genes, potentially enhancing adaptability and pathogenicity [16, 34]. These findings highlight the importance of recombination-aware genomic surveillance to monitor emerging variants and inform public health responses.

The most significant virulence factor of *C. diphtheriae* is diphtheria toxin, a highly potent exotoxin mediated via the *tox* gene carried in some corynebacteriophages [1]. PCR methods and genomic approaches have been honed to predict the presence of the *tox* gene accurately. This study demonstrated complete concordance between genomic and PCR methods for detecting the *tox* gene in contemporary isolates, providing confidence that genomic surveillance alone can adequately detect *tox* gene presence, at least in our setting. Phenotypic expression of the *tox* gene was not assessed using the Elek test [36] in this study, as the required antitoxin is not available in Australia. However, genomic analysis revealed no evidence of non-functional *tox* genes, suggesting the absence of NTTB strains in this dataset. Our phylogenetic analysis revealed that *tox* gene positive isolates were dispersed in multiple unrelated sublineages, suggesting independent acquisition of the *tox* gene as observed in previous studies [16, 24]. Nevertheless, events involving the acquisition or loss of the *tox* gene within the same lineage have been previously reported in *C. diphtheriae* [16, 24, 33], and similar findings were observed in this study, with both *tox* positive and *tox* negative isolates of ST20 and ST124 identified (Figure 1, Supplementary Table S1).

This study identified a predominantly *tox* positive population among historical *C. diphtheriae* strains and a predominantly *tox* negative population in contemporary strains in Victoria. This finding reflects the long-term impact of widespread immunisation against diphtheria in the region, contributing to genomic and population-level shifts in *C. diphtheriae* strains in Victoria. The observed pattern suggests an evolutionary trajectory away from toxin production, potentially driven by vaccine-induced host immunity, altered transmission dynamics, and reduced fitness of toxigenic strains in a predominantly immunised population [37]. These findings emphasise the importance of incorporating historical *C. diphtheriae* genomes into genomic analyses, as they could provide crucial information on strain turnover and shifts in lineage distribution in the population. Such insights are critical for understanding pathogen population dynamics over time and for guiding ongoing genomic surveillance efforts.

Recent re-emergence of diphtheria has posed a public health burden to some countries [22, 38–40]. Comparatively, diphtheria outbreaks are extremely rare in Australia and no diphtheria outbreaks have been reported in Victoria in the last two decades. The most recent outbreak of *tox* positive *C. diphtheriae* (of ST381) occurred in North Queensland, Australia in 2022 [4], with 24 locally acquired cases reported between January and August. Most isolates (n=18) were from cutaneous samples, while six were respiratory, although only two patients exhibited classic respiratory diphtheria symptoms. No mortalities were reported. There was no evidence of transmission to Victoria and ST381 was not observed among Victorian isolates in this study. However, in the event of re-emergence of diphtheria in Australia, genomics will play a critical role in identifying likely transmission routes, distinguishing multiple importations from occult local transmission. Through this study, we have established the genomic baseline and methods to allow these data to be rapidly produced and analysed, generating actionable data for public health responses.

Analysis of *tox* negative contemporary isolates in this study revealed a high prevalence of ST32, which were epidemiologically not related. Apart from Australia, ST32 has been reported in European countries [13, 24, 33]. *C. diphtheriae* strains of ST32 were observed to have enhanced virulence due to increased adhesion potential [13], possibly contributing to the relative frequency in the Australian population. ST32 was reported in the state of Victoria (this study) and New South Wales [13], but ST39 has been frequently reported among *tox* negative isolates in North Queensland [4], demonstrating some genetic diversity of non-toxigenic *C. diphtheriae* strains between geographic locations within Australia.

Emergence of antimicrobial resistance in *C. diphtheriae* poses a significant challenge to the clinical management of *C. diphtheriae* infections, as well as to control pathogen transmission. Unfortunately, clinical susceptibility breakpoints lack standardisation between the two dominant susceptibility testing guidelines, EUCAST and CLSI, posing challenges in detection and comparison of AMR in a global context [16, 33]. Some studies have proposed implementing tentative epidemiological cut-off (TECOFF) values for antimicrobials [16, 41], however, no TECOFF values are currently publicly available for any antimicrobial during this study. Whilst discordances between clinical breakpoints remain, this study at least provides some baseline AMR data for interpretation and to potentially guide empiric therapy.

While previous studies have compared resistant phenotypes to genetic determinants of AMR in *C. diphtheriae*, very few Australian isolates were investigated in such studies [16, 19, 42]. In this study, we provide a comprehensive overview of both antimicrobial resistant phenotypes as well as known genetic determinants of AMR in *C. diphtheriae* isolates from Victoria, Australia. Our analysis of phenotypic resistance against first-line antimicrobials demonstrated an emerging resistance to penicillin (17.7%) and erythromycin (5.6%) in contemporary isolates compared to no resistance to both penicillin and erythromycin among historical isolates. Despite following different susceptibility testing guidelines, increased penicillin resistance or reduced susceptibility rates have been reported in multiple studies [16, 42, 43]. In contrast, phenotypic resistance or reduced susceptibility to erythromycin remains relatively low [16, 22, 43]. Phylogenetic distribution of antimicrobial resistance genes showed their presence across multiple unrelated sublineages (Figure 1), indicating independent acquisition via horizontal gene transfer, consistent with previous studies [16, 33].

In conclusion, we have used comprehensive genomic analysis to demonstrate a highly diverse population structure in Victorian *C. diphtheriae* in both historical (predominantly *tox* positive) and contemporary (predominantly *tox* negative) strains, establishing a baseline for future investigations. The Victorian sequences are dispersed across the global *C. diphtheriae* phylogeny, reflecting multiple independent importations in the contemporary setting. We have demonstrated high concordance between PCR and *in silico* methods for detection of diphtheria toxin gene (*tox*) and shown the added value of genomic AMR detection to augment traditional phenotypic AST methods. Given the potential for global re-emergence of diphtheria, particularly in areas of conflict and low vaccine coverage and with the added complication of increasing AMR, it is critical for public health and clinical laboratories in all settings to prepare for future epidemics of *C. diphtheriae*. As we have demonstrated, genomic surveillance likely plays a significant role in identifying transmission routes and patterns to inform public health actions. Future work should focus on providing equitable access to genomic and phenotypic surveillance across all countries, including low– and middle-income countries, enabling effective global surveillance and coordinated responses.

## Author statements

### Author contributions

L.S.K. – conceptualization, investigation, data curation, analysis and data visualisation, writing original draft. S.B. – data analysis, writing-review and editing. K.H. – investigation, writing-review and editing.

J.S. – data curation, writing-review and editing. N.L.S. – conceptualization, writing-review and editing.

B.P.H. – conceptualization, writing-review and editing.

## Conflicts of interest

The authors declare that there are no conflicts of interest.

## Funding information

The Microbiological Diagnostic Unit Public Health Laboratory is funded by the State Government of Victoria. N.L.S and B.P.H. are supported by NHMRC Investigator Grants (GNT2033803 to N.L.S., GNT1196103 to B.P.H.).

## Ethical approval

All samples were collected as part of routine public health laboratory surveillance, under the Public Health and Wellbeing Act 2008 (Victoria), and under previous equivalent public health legislation (e.g. Public Health Act 1958). Ethical approval for this study was received from the University of Melbourne Human Research Ethics Committee (study number 1954615.3).

## Acknowledgements

We acknowledge and thank the diagnostic laboratories in the state of Victoria for referring *C. diphtheriae* isolates to Microbiological Diagnostic Unit Public Health Laboratory (MDU PHL) and supporting the surveillance of *C. diphtheriae* in Victoria. The authors gratefully acknowledge Ms Kerrie Stevens for her expert laboratory assistance and all MDU PHL staff who involved in processing, phenotypic and molecular characterisation and sequencing of the samples used in this study.

## Abbreviations

AMR: antimicrobial resistance
*tox*: diphtheria toxin gene
NTTB: non-toxigenic *tox* gene-bearing
MDU PHL: Microbiological Diagnostic Unit Public Health Laboratory
PCR: polymerase chain reaction
MLST: multi-locus sequence type
ST: sequence type
MIC: minimum inhibitory concentration
MDR: multidrug-resistant
CLSI: Clinical and Laboratory Standards Institute
EUCAST: European Committee on Antimicrobial Susceptibility Testing

## Notes

### Competing Interest Statement

The authors have declared no competing interest.

